# A single-cell transcriptomic atlas of inner ear morphogenesis in zebrafish

**DOI:** 10.1101/2025.09.30.679639

**Authors:** Akankshi Munjal, Kalki Kukreja, Samara Williams, Toru Kawanishi, Natasha M. O’Brown, Kana Ishimatsu, Allon Klein, Sean G. Tsung-Megason, Ian A. Swinburne

## Abstract

The inner ear constitutes different cell types next to one another: the sensory patches whose hair cells synapse with neurons, the thin channels of three semicircular canals whose perpendicular organization enables detection of directional head rotation, and the endolymphatic duct and sac whose conditional epithelial barrier relieves excess pressure and promotes fluid pressure homeostasis. How the ear’s component cell states are established during development has remained unknown. We use single-cell RNA sequencing to distinguish cell states within the developing ear with wild-type zebrafish embryos and *lmx1bb* mutants that exhibit defects in canal and sac morphogenesis. We identify the earliest marker for the semicircular canal-genesis zone (*ccn1l1*), unexpected genes in the endolymphatic sac that suggest a role for tissue contraction in its function (*smtnb*), parallel gene sets for sensory patches in the neuromast and ear, and a conserved role for cell-cycle pausing (*cdkn1bb* expression in the canals and sac as previously observed in the developing mouse ear). This atlas provides the most comprehensive transcriptional profiling of the developing inner ear, identifying new molecular leads to understand ear morphogenesis.

**Summary statement:** Single-cell transcriptomic analysis of developing wild-type and *lmx1bb* mutant zebrafish reveals cell-states and effectors that distinguish the inner ear’s sensory patches, semicircular canals, endolymphatic duct and sac, and periotic mesenchyme.

## Introduction

The two primary functions of the inner ear are to detect sound and maintain balance. The adult zebrafish inner ear is composed of connected chambers: the lagena that senses sound vibrations (closest to the cochlea), the three semicircular canals that detect head rotations, and the endolymphatic duct and sac that maintain a stable composition of the pressurized fluid called endolymph encased in these chambers. A fundamental question concerning inner ear function is how different cell types develop. This question can be broken into three parts: what are the molecules that distinguish the different cell states of the inner ear during development; how do different cell states organize in space over time; and which biochemical networks initiate and maintain inner ear function. We address the first of these questions by profiling the transcriptome of the developing inner ear of zebrafish using single-cell mRNA sequencing (scRNA-seq). We use the optical and physical accessibility of the zebrafish inner ear to provide insights into spatial organization with whole-mount multiplex fluorescent in situ hybridization (mFISH) imaging of the predicted cell states emerging from scRNA-seq. Advancements in scRNA-seq combined with mFISH allow for the systematic mapping of gene expression to create “cell state atlases”, building an essential foundation for informed hypotheses and functional analysis.

The hair cells are the primary cell type that allow the inner ear to detect sound and head movement for balance. Hair cells convert mechanical stimuli from fluid movements, which bend their thin protrusions, into bioelectrical signals through a process called mechanotransduction(Hudspeth and Corey 1977; Teresa Nicolson 2017; Ó Maoiléidigh and Ricci 2019; Shotwell et al. 1981). The distinctive and critical role of hair cell function for hearing and balance has led to the identification of proteins and activities critical for hair cell physiology through the application of genetics to the study of both model organisms and human deafness(T. B. Friedman and Griffith 2003; L. M. Friedman et al. 2007; Ingham et al. 2019; Teresa Nicolson 2017; T. Nicolson et al. 1998; Whitfield 2002).

In the adult zebrafish inner ear, the sensory patches known as cristae— composed of hair cells and supporting cells located at the base of each semicircular canal—detect angular acceleration during head rotation (Fig. 1A, dark blue). The otolithic organs (utricle, saccule, and lagena) contain sensory patches called maculae that detect linear acceleration and gravity, with the saccule and lagena also contributing to auditory function. A recent study profiled zebrafish inner ear hair cells and identified two subtypes that share gene expression patterns with mammalian hair cells, suggesting common ancestry (Shi et al. 2023). In addition to the hair cells within the inner ear, fish also have patches of hair cells called neuromasts (NMs) in their skin concentrated in an organ called the lateral line (Fig. 1D, dark green). Because of their superficial nature, ability to regenerate, and similarity to the ear’s hair cells, studies of the NM have produced a wealth of knowledge regarding their molecular genetics. We sought to determine which transcriptional programs contribute to the development of the ear’s hair cells and how they might differ from the better-studied hair cells of NMs (Lush et al. 2019).

**Fig. 1.**
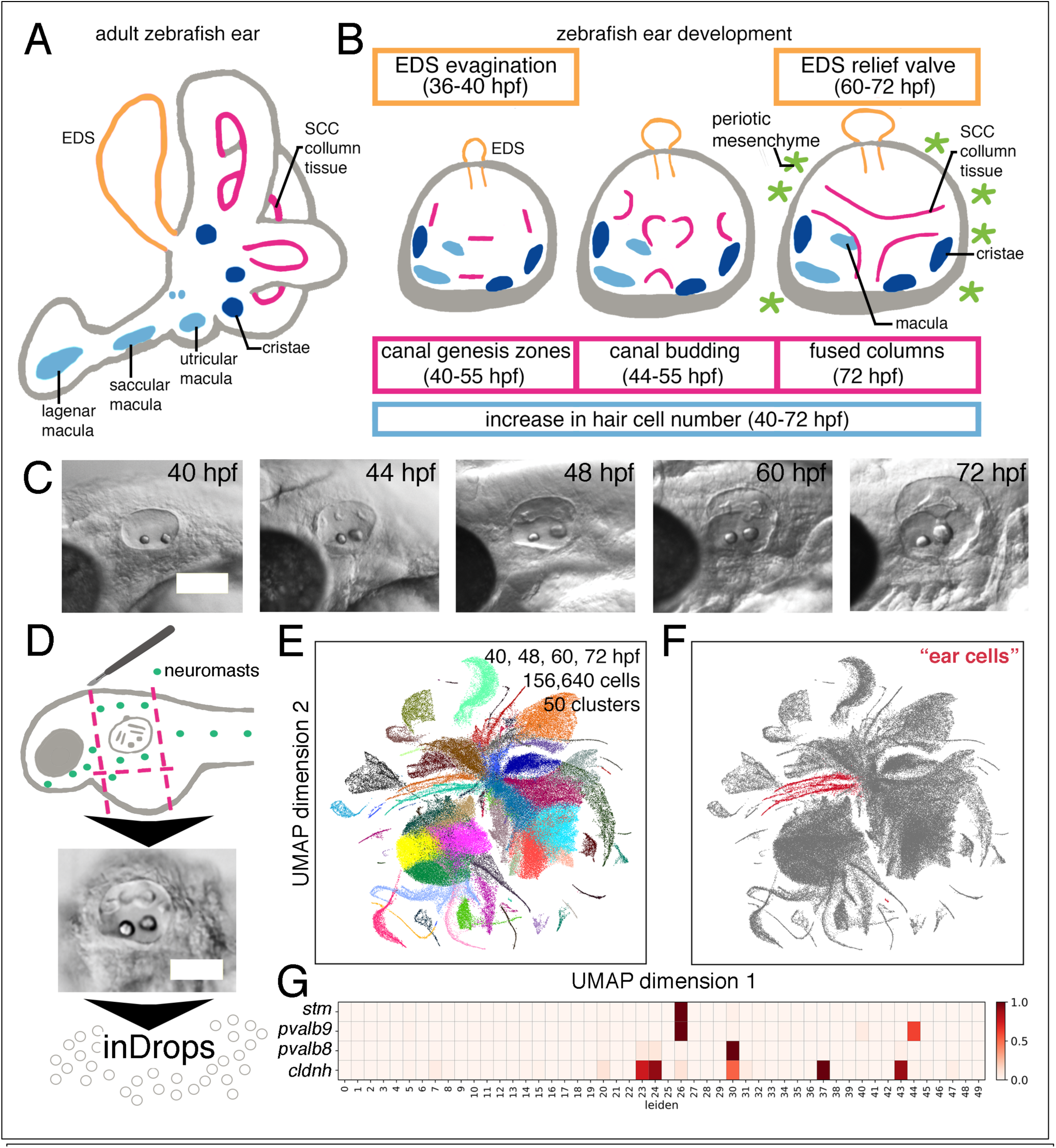
Coarse mapping of single-cell transcriptomes for zebrafish cells enriched for the ear. A. Cartoon schematic of ear anatomy that highlights tissue and cell types in the adult zebrafish ear. B. Cartoon schematic of ear developmental events queried by our experiments. C. Brightfield images of representative developmental stages examined. D. Cartoon schematic of experimental workflow. E. UMAP graph of cells clustered by transcriptomes. F. UMAP graph with presumptive ear and neuromast cells in red. G. Heat map of normalized expression indicating clusters that have cells expressing known markers of the ear and neuromast. (Scale bars in C, D 50 μm)

The function and the survival of the delicate hair cells require a stable composition of endolymph fluid. The endolymphatic duct is a narrow channel that connects the endolymphatic sac (ES), a dead-end chamber, to the fluid within the rest of the inner ear (Fib. 1A-B, orange). The ES was thought to have roles in stabilizing a fluid composition necessary to consistently sense sound for hearing and movement for balance(Naito 1950; Salt 2001). We previously found that the lumen of the ES periodically inflates and deflates like a balloon (Swinburne et al. 2018), behaving like a relief valve through the periodic opening of junctions to maintain a consistent volume and pressure within the ear. We sought to identify transcriptomic changes underlying the developmental transition from the naive dorsal epithelium into the conditional epithelial barrier of the endolymphatic duct and sac.

The embryonic structure that gives rise to the adult zebrafish inner ear is an ellipsoidal epithelial shell called the otic vesicle (OV) that emerges from the cavitation of the ectodermally derived otic placode(Alsina and Whitfield 2017; C. Haddon and Lewis 1996; Mosaliganti et al. 2019; Whitfield 2002). For the OV to attain the ability to detect rotational acceleration, it undergoes dramatic topological changes. During zebrafish development, semicircular canal morphogenesis begins when six discrete groupings of epithelial cells bud inward from the wall of the OV and grow toward one another driven by force generated from the secretion of extracellular matrix (magenta, Fig. 1B-C)(Munjal et al. 2021; C. M. Haddon and Lewis 1991). Sequentially, buds grow from the lateral wall, the anterior wall, the posterior wall, and then finally from the ventral wall of the OV. Except for the ventral bud, all canal buds emerge roughly at the midpoint of the D-V axis. These buds then grow inwards towards one another driven by force generated from the secretion of the extracellular matrix(C. M. Haddon and Lewis 1991; Munjal et al. 2021). Pairs of projections meet and fuse to form three columns, thereby demarcating the three canals. BMP, FGF, RA, SHH, and Wnt signaling pathways have been implicated in the initial patterning of the OV but it is unknown how these signals are integrated to first pattern the canals(Chang et al. 2004; Choo et al. 1998; Gerlach et al. 2000; Mansour et al. 1993; Noda et al. 2012; Pauley et al. 2003; Rakowiecki and Epstein 2013; Riccomagno et al. 2005). Recently, the transcription factor Lmx1b has been shown to be involved in promoting extracellular matrix synthesis genes at the buds(Mori et al. 2025). However, we do not yet know what cell state first distinguishes the ‘canal genesis zones’(Chang et al. 2004). Following canal budding, we do not know the global collection of gene expression states and molecular activities that contribute to the sculpting of the semicircular canals.

Classic embryological experiments suggested back and forth communication between the developing epithelium of the OV and the surrounding “periotic” mesenchyme (Fig. 1B, light green). The outcome of this communication is that signals from the developing ear can induce the mesenchyme to become a cartilaginous capsule. Conversely, signals from the periotic mesenchyme contribute to the development of the ear’s semicircular canals and sensory tissues (Kaan 1930, 1938). To identify new leads to the molecular basis of how these two tissues induce and influence one another, we sought the transcriptomic distinctions of the periotic mesenchyme’s cell-state during a developmental period that appears to be the initiation of capsule condensation.

Zebrafish mutants that lack the Lmx1bb transcription factor develop differently with extra hair cells, incomplete morphogenesis of semicircular canals, and an endolymphatic sac that cannot relieve excess luminal pressure (Mori et al. 2025; Swinburne et al. 2018; Obholzer et al. 2012; Schibler and Malicki 2007). We used single-cell RNAseq of developing wild-type and *lmx1bb* mutant to generate gene lists that may contribute to the reported phenotypes.

Within our gene expression atlas we find states including new molecular candidates for uncovering the mechanistic answers to several mysteries of ear development. Given the combined importance of the ear’s geometry for its function and requirement for control of mechanical forces from hydrostatic pressure and ECM production to sculpt its tissues and maintain homeostasis, these new molecular handles will be instrumental for future studies of how information flows between biochemical, mechanical, and geometric modes for morphogenesis(Collinet and Lecuit 2021).

## Results

### Identifying ear cell states by unsupervised clustering of single-cell transcriptomes

The data set we report here consists of 156,640 single cell transcriptomes spanning multiple time points of development and encompassing the developing ear. To generate this data, we forwent sorting of cells so as not to overlook any cell-types. Instead, to enrich for the otic epithelial tissue, we dissected embryos to remove as much of the residual embryo as possible (Fig. 1D). We collected wild-type samples at 40, 48, 60, and 72 hours post fertilization (hpf) because these developmental time-points span the patterning and morphogenesis of the semicircular canals (SCCs), the endolymphatic duct and sac (EDS), and maturation of hair cells (Fig. 1B-C). We also collected *lmx1bb* mutant samples at 60 and 72 hpf because these developmental time-points correspond to early developmental phenotypes with apparent delayed budding of the SCCs and absence of relief valve activity in the ES(Obholzer et al. 2012; Swinburne et al. 2018). In total, we dissected approximately 600 fish and sequenced the barcoded transcriptomes using inDrops scRNA-seq (Fig. 1D). After sequencing, data was embedded for clustering and visualization following standard approaches to detect highly variable genes, reduce dimensionality by principal component analysis (PCA), and the Leiden algorithm used to partition the cells into 50 clusters (Fig. 1E). Two of these clusters corresponded to crude admixtures of cells of the OV and NM (Fig. 1F) as determined by differential expression of marker genes, which included *stm* and *pvalb9* for the OV and *cldnh* and *pvalb8* for the NM (clusters 26 and 30, Fig. 1G). The mixing was expected due to the close similarity of hair cells in the two organs. Other clusters include diverse cell types, particularly from different brain regions and the skin, which we have not analyzed further here. Supplemental Table 1 lists statistically significant enriched genes for all 50 clusters and our annotations of the remaining clusters. Processed transcriptome data has been deposited in Zenodo, 10.5281/zenodo.17188954; and the raw ScRNA-seq data have been made available on Gene Expression Omnibus with accession number GSE309286.

### Coarse map of partitioned cells of the ear and neuromast

To begin to characterize cell-states within the ear, we combined and further partitioned the otic vesicle (OV) and neuromast (NM) clusters using the same unsupervised clustering pipeline into 13 different clusters (Figs. 1F, 2A). We identified genes that were enriched in each cluster and used these to annotate clusters corresponding to different populations of cells, for which we then calculated and graphed the relationships of populations of cells using hierarchical clustering (Fig. 2A, D). We used the expression of known OV and NM genes to determine which clusters belong to the respective organs (*pvalb8, pvalb9, stm,* Fig. 2B). We identified and annotated NM clusters as those that included cells of the NM hair cell lineage (3,4) based on expression of *dlb* and *pvalb8* and supporting cells of the NM (5, 6, 7) based on the expression of genes like *sostdc1a, fat1b, hopx,* and *krt4* (Fig 2 A,D, Supplemental Table 2) (Lush et al., 2019). We annotated OV clusters as those that included less differentiated epithelial cells (clusters 12, 13) based on the respective enriched expression of genes like *emilin1a* and *bricd5*, cells of the OV hair cell lineage (1,2) based on enriched expression of genes like *pvalb9* and *atoh1a*, cells of the SCC tissue (9) based on enriched expression of genes like *vcanb* and *matn4*, and cells of the EDS (10) based on enriched expression of genes including *foxi1* and *wnt16* (Supplemental Table 2 lists genes with statistically significant enriched expression). Altogether, our pipeline to enrich the ear cell population, unsupervised clustering, and manual annotation resulted in 3363 annotated OV and NM cells, facilitating statistical analysis of differentially expressed genes.

**Fig. 2.**
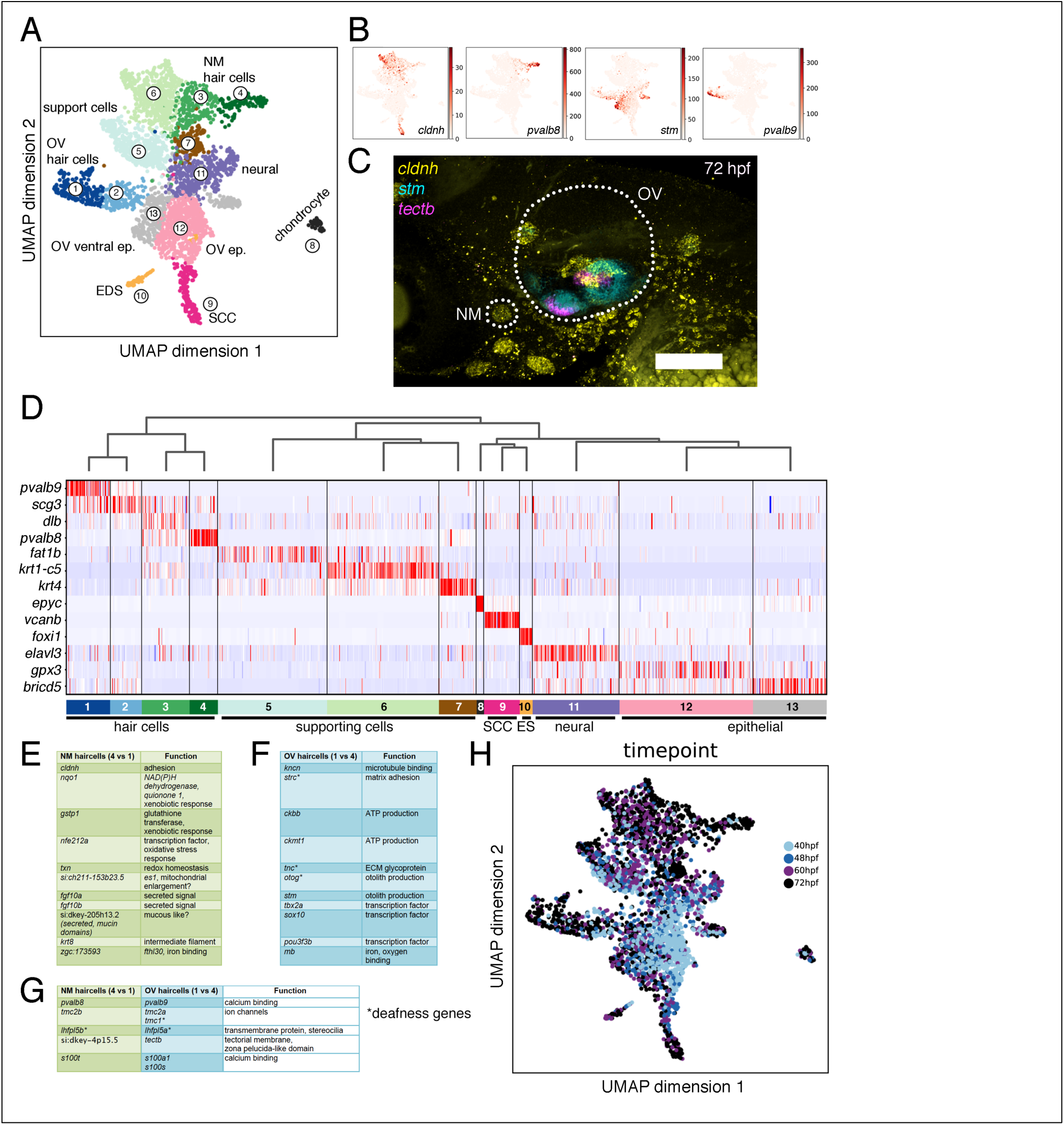
Identifying ear and neuromast cell states by unsupervised clustering of transcriptomes. A. UMAP plot of clustered ear and neoromast cells based on single-cell transcriptomes. B. Heat maps of transcript counts for expression of nueromast and ear markers overlayed on UMAP graph from A. C. Multiplexed in situ hybridization analysis of ear and neuromast markers. D. Hierarchical organization of clusters form A. with heat map histogram of markers for each cluster. E. Examples of genes enriched in expression in neuromast hair cells relative to otic vesicle hair cells. F. Examples of genes enriched in expression in otic vesicle hair cells relative to neuromast hair cells. G. Examples of genes differentially enrich between otic vesicle hair cells and neuromast hair cells that appear to either be paralogs or perform homologous roles. H. Color-coding of developmental stage on UMAP plot from A. (Scale bars in C 50 μm)

*cldnh* is expressed in the neuromast and OV (Gong et al. 2021), while *stm* and *tectb* are both elevated in cells of the ventral OV but are known for distinct roles in otolith formation(Söllner et al. 2003) versus otolith embedding in the tectorial membrane(Stooke-Vaughan et al. 2015). To determine whether there is a spatial distinction in the cells expressing the markers identified, we used the multiplexed and sensitive fluorescent in situ hybridization technique, Hybridization Chain Reaction (HCR)(Choi et al. 2018a), to visualize the expression of *cldnh*, *tectb,* and *stm* (yellow, magenta, and cyan, Fig. 2C) and confirmed that *stm* and *tectb* are primarily expressed in a subset of cells within the OV and *cldnh* is primarily expressed in NM (except for expression within the columns of the SCC during morphogenesis). With expression of markers like *pvalb8* and *cldnh* of NM and *pvalb9* and *stm* of OV, the clusters organized into NM related cells at the top of the map and OV related cells at the bottom of the map (Fig. 2A)(Thisse and Thisse, C. 2004). To dissect distinctions between OV and NM hair cells, we compared enriched genes between populations of hair cells and uncovered differences (e.g., fgf10a/b, sox10), as well as differential use of paralogs in the two populations (Fig. 2E-G).

In addition to the expected cell types associated with the OV and NM, we found ear related “neural” cells (11) based on expression of markers like *elavl3* and “chondrocyte-like” cells (8) based on expression of markers like *epyc.* Neurons that innervate the hair cells of the OV originate from the ventral otic epithelium. To address the identity of these ear related neural cells, we compared their transcriptional profiles to those of all other neural cells in our dataset (clusters 1, 3, 5, 6, 12, 14, 16, 34, and 41 of Fig. 1E and Table 1). The primary difference of the ear related neurons relative to other neurons was the expression of ear and neuromast markers such as *stm.* In addition to these ear markers, the ear related neural cells were distinguished from all other neural cells by expression of genes implicated in neuroblast development from the ventral otic epithelium, such as *eya1*, *jag1b, hey1*, *spry4*, *epha7*, *sox2*, *sema3fb*, and *gata3* (Supplemental Table 4) (Ahmed et al. 2012; Gou et al. 2018; Kantarci et al. 2015; Karis et al. 2001; Kozlowski et al. 2005; Pan et al. 2010; Puligilla and Kelley 2017; Puligilla et al. 2010; Scott et al. 2019; Vemaraju et al. 2012).

### Fine map and paths of transcriptional states for hair cells and supporting cells

The distribution of cells from progressive developmental times within the UMAP plot of partitioned OV and NM cells (Fig. 2H) suggest that the sampled cell states order along a developmental continuum. This continuum spanned between cells from 40 and 48 hpf that are enriched in the least-differentiated clusters like the OV epithelium (12, 13) and the NM support cells (5,6), to cells from 60 and 72 hpf that are enriched in more-differentiated clusters, including the OV hair cells, SCC, EDS, and the NM hair cells (1, 9, 10, 4).

We sought to improve the interpretability of these features within the data by determining the distance relationships for separate lineages (hair cells, SCC, or ES). During the homologous developmental time window, the mammalian ear shows programmed senescence, marked by downregulation of cell cycle genes and induction of cell cycle inhibitors, consistent with the minimal cell division observed in zebrafish during these developmental windows(Muñoz-Espín et al. 2013; Swinburne et al. 2018; Munjal et al. 2021). Furthermore, because regions mature at different times, as in the staggered morphogenesis of semicircular canals, we applied pseudotime analysis to trace cell trajectories. To refine the lineage-specific maps, we first repartitioned relevant subsets of clusters from the full OV/NM set and then connected the cell-states by distance relationships using partition-based graph abstraction (PAGA, Fig. 3A, B)(Wolf et al. 2019). The organization of cells in the PAGA-graph reflected their position in developmental time (Fig. 3C). The NM cell states resemble those previously described at 5 days post fertilization; however, some populations appear to be less distinctive, likely reflecting differences between developmental and homeostatic programs (graphed heatmaps of expression for representative markers, Fig. 3D, Supplemental Table 4, (Lush et al. 2019)). At the earlier time points we studied (40-72hpf), mantle-like cells express genes such as *fat1b* and *sfrp1a,* which are mantle cell markers at 5 dpf (cluster “n0”, unique cluster identifier where letter indicates which map, i.e. the map in Fig.3 uses “n”) (Lush et al., 2019). Clusters n1-3 represent populations of proliferating progenitors (*pcna*+ and *bmp5a*+) distinguished by differential expression of genes like *cyp26c1* (n1), *pvalb2* (n2), and *krt4* (n3). Further along this path are immature hair cell populations expressing delta genes (*dlb*, *dld*) and *atoh1a* (n4) and mature hair cells expressing physiological genes like *pvalb8* (n5). Appearing to diverge from the NM hair cell path are two populations of NM supporting cells (n7 and n8). One of the strongest markers of n7 is an uncharacterized zebrafish gene *si:dkey-4p15.5.* InterProScan predicts that the protein product of *si:dkey-4p15.5* is secreted with a N- and C-terminal zona pelucida domain, similar to proteins of the ear’s tectorial membrane proteins, such as Tectb (44% pairwise identity)(Blum et al. 2025). Cluster n8 resembles the ring cells of the 5 dpf NM with the expression of ring markers like *hopx* (Lush et al., 2019). However, cluster n8 is also distinguished by the expression of 5 dpf mantle markers such as *ponzr6* and *tspan1* (Lush et al. 2019). A heatmap of differential gene expression and distance in gene expression space summarizes paths of the NM population, including expression of proliferation marker (*pcna*) and “G0” marker (*cdkn1bb*) (Fig. 3E).

**Fig. 3.**
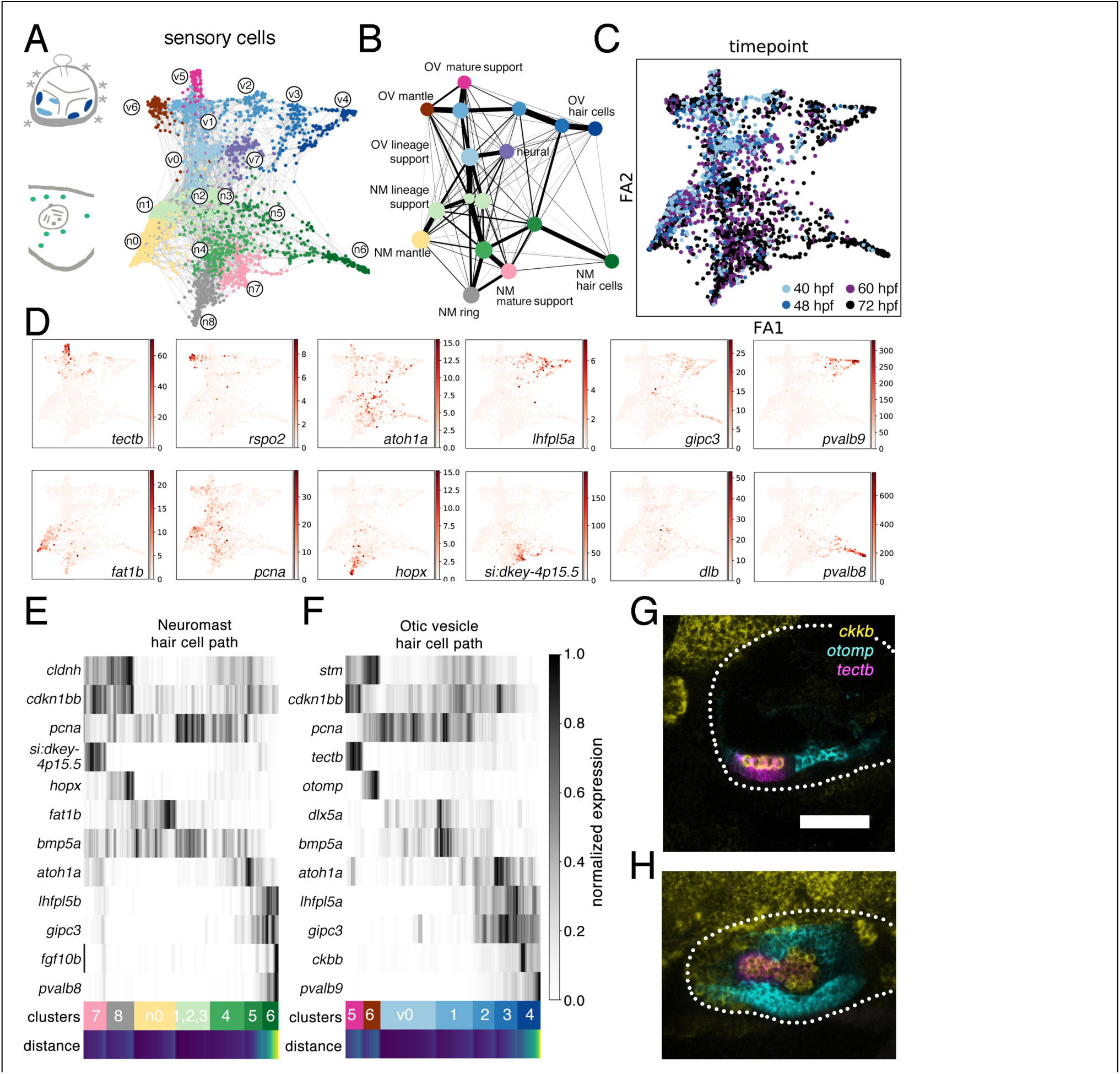
Fine analysis for single-cell transcriptomes of hair cell paths in the neuromast and the otic vesicle. A. Cartoon schematics of a 3 day old zebrafish ear, with otic vesicle hair cells highlighted in blue, and a cartoon of neuromasts with hair cells highlighted in green next to a PAGA organized graph of hair cells. B. Coarse-grained PAGA graph of hair cells. C. Color-coding of developmental stage on PAGA graph. D. Heat-map of transcript counts on PAGA graph with markers representative of different clusters. E. Collapsed heat-map histogram of normalized expression that compares gene expression in cells along the PAGA path of neuromast hair cells with calculated distance along the path. F. Collapsed heat-map histogram that compares gene expression in cells along the PAGA path of otic vesicle hair cells with calculated distance along the path. G, H. Multiplexed fluorescent in situ analysis of otic vesicle genes of interest from the PAGA graph. (Scale bar 50 μm, same for G,H)

The path of the OV sensory cells echoes that of the NM beginning with populations of proliferating neuroepithelial progenitors (v0, v1). Expression of marker genes such as *lhfpl5a*, *gipc3*, and *atoh1a* in clusters v2 and v3 indicates that these populations represent immature hair cells. The path continues to cluster v4 where the expression of physiological markers like *pvalb9* reflects the molecular state of more mature hair cells. Like the NM path, the otic sensory path has populations of cells that branch away from the hair cell lineage. Cluster v5 represents a population of support cells that produce the tectorial membrane protein Tectb. Cluster v6 is a distinct population of supporting cells characterized by the expression of *otomp* and genes encoding Wnt signaling modulators such as *rspo2* and *sfrp5*, like the mantle cells of the NM (cluster n0, *sostdc1a* and *sfrp1a*). A heatmap of marker gene expression and distance in gene expression space summarizes paths of the OV ventral neuroepithelium and sensory cells (Fig. 3F).

To confirm whether the distinct cell-states identified by scRNAseq reflect spatially distinct patterns within the developing otic vesicle, we performed HCR fluorescent in situs (Fig 3G-H). At 60 hpf, expression of support cell marker *tectb* (magenta, representative of cluster v5) flanks hair cells expressing *ckbb* (yellow, representative of cluster v4), which are both surrounded by a population of support cells expressing *otomp* (cyan, representative of cluster v6).

### Fine map and path of transcriptional states for cells that shape the semicircular canals

The semicircular canals form from the cells in the dorsal otic epithelium. To refine the map of cell-states that contribute to semicircular canal formation, we pooled cells from the non-ventral otic epithelium and SCC clusters (clusters 12 and 9, Fig 4A, supplemental table 2), repartitioned these cells, calculated their distance relationships, and then plotted the PAGA-initialized embedding (Fig. 4A) and coarse-grained PAGA-graph (Fig. 4B). The organization of cells in the PAGA-graph reflected their position in developmental time (Fig. 4C). The path begins with cell states in clusters c1-5 characterized by the expression of proliferation markers (*pcna, hmgb2a*, markers for all clusters in Supplemental Table 5). Cluster c6 lacks significant gene markers but appears to represent a cell state in transition toward the onset of morphogenesis with low expression of genes like *lmx1bb,* known to be involved in the process. Clusters c7-9 appear to encompass cells involved in column budding, growth, and fusion and are distinguished by expression of *jun* (c7), *serpinh1b* (c8), and *nr4a3* (c9), in addition to sharing expression of known SCC genes like *ugdh* and *vcanb* (Fig. 4D). A heatmap of marker gene expression and distance in gene expression space summarizes paths of cell states underlying SCC morphogenesis (Fig. 4E).

**Fig. 4.**
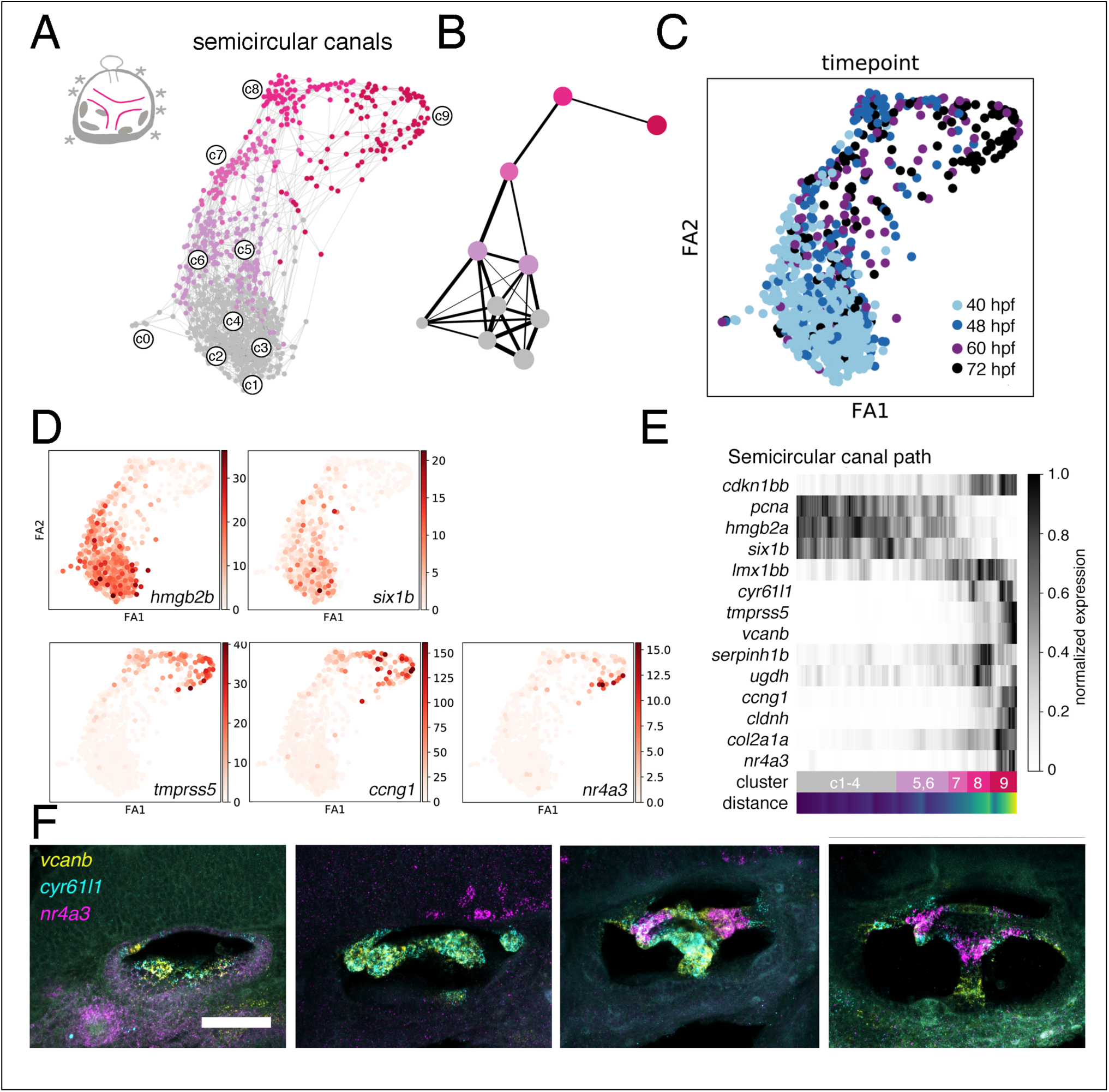
Fine analysis for single-cell transcriptomes of semicircular canal cells. A. Cartoon schematic of a 3 day old zebrafish ear, with canal cells highlighted in magenta, next to PAGA organized graph of SCC cells with more “mature” SCC cells colored magenta. B. Coarse-grained PAGA graph of SCC cells. C. Color-coding of developmental stage on PAGA graph. D. Heat-map of transcript counts on PAGA graph with markers representative of different clusters. E. Collapsed heat-map histogram that compares gene expression in cells along the PAGA path with calculated distance along the path. F. Multiplexed fluorescent in situ analysis of genes of interest from the PAGA graph. (Scale bar 50 μm)

Intrigued by salt-and-pepper expression of some SCC specific genes, we used multiplexed HCR in situ hybridization to visualize the expression of *vcanb, ccn1l1* (also known as *cyr61l1*), and *nr4a3* (Fig. 4F). We found two unexpected patterns. First, we found that *ccn1l1* is expressed in a restricted manner to a patch of cells, coincident with the presumptive bud sites or the “canal-genesis zone”, 2-4 hours before the epithelium thickens and buds out. This pattern represents the earliest known SCC specific expression pattern and provides a handle for future studies that will seek to understand how the SCCs are robustly patterned. The second unexpected pattern concerns *nr4a3,* whose expression does not appear to turn on until two SCC columns meet. We speculate that *nr4a3* may be part of an expression program for later stages of canal morphogenesis(Ponnio et al. 2002).

### Fine map and path of transcriptional states for cells in the endolymphatic duct and sac

Like the semicircular canals, the endolymphatic duct and sac (EDS) also develops from the dorsal OV. To refine the map of cell-states that contribute to the endolymphatic duct and sac (EDS), we pooled cells from the non-ventral otic epithelium and the annotated EDS clusters (12 and 10, Fig 2A), repartitioned these cells, calculated their distance relationships, and then plotted the PAGA-initialized embedding (Fig. 5A) and coarse-grained PAGA-graphs (Fig. 5B). Again, the organization of cells in the PAGA-graph reflected their position in developmental time (Fig. 5C). The path begins with cell states in clusters e0-4 characterized by the expression of proliferation markers (*pcna, hmgb2a*, subset of markers in Fig. 5D, markers for all clusters of Fig. 5A in Supplemental Table 6). Cluster e6 is characterized by the expression of the marker *apcdd1l*. Cluster e7 appears to include the cells of the EDS with expression of known markers like *foxi1*. We identified 126 genes significantly enriched for expression in the EDS relative to the rest of the otic epithelium (Supplemental Table 6). A heatmap of marker gene expression and distance in gene expression space summarizes paths from dorsal otic epithelium to the cells of the EDS (Fig. 5E). We used HCR to visualize the expression of EDS markers *foxi1, wnt16,* and *smtnb* (Fig. 5F-H). These findings indicate that there are clear transcriptional differences between the cells of the duct (*wnt16,* proximal to the otic vesicle) and cells of the sac (*smtnb,* most distant from the interior of the otic vesicle). The expression of *smoothelin b*, normally thought to be limited to smooth muscle cells, suggests regulated sac cell contraction.

**Fig. 5.**
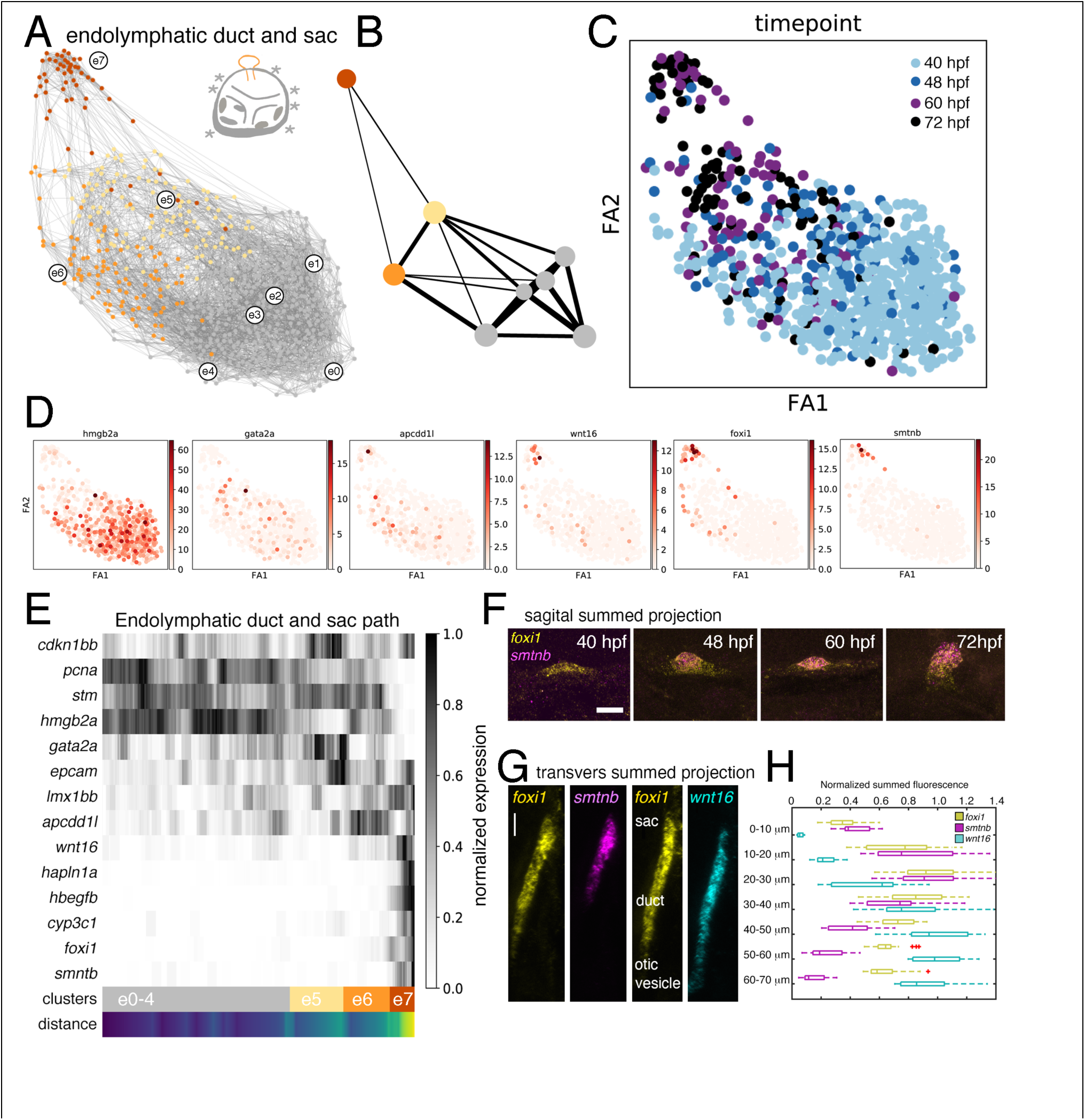
Fine analysis for single-cell transcriptomes of endolymphatic duct and sac cells. A. Cartoon schematic of a 3 day old zebrafish ear, with the endolymphatic duct and sac highlighted in orange next to PAGA organized graph of dorsal otic vesicle and EDS cells with more “mature” EDS cells colored orange. B. Coarse-grained PAGA graph of EDS cells. C. Color-coding of developmental stage on PAGA graph from A. D. Heat-map of PAGA graph with markers representative of different clusters. E. Collapsed heat-map histogram that compares gene expression in cells along the PAGA path with calculated distance along the path. F. A sagittal perspective of multiplexed fluorescent in situ analysis of EDS genes expressed at the queried developmental stages. G. A transverse perspective of multiplexed fluorescent in situ analysis of duct and sac genes expressed at 72 hpf. H. Quantification of the in situs represented in G. (Scale bars 10 μm)

### Fine map and path of transcriptional states for cells of the periotic mesenchyme

Cells of the OV and NM clusters express orthologs of many known human deafness genes (Supplemental Figure 1). When we reassessed the other clusters in our full dataset, cluster 28 also stood out for the statistically significant expression of genes for which mutations in the human orthologs can cause deafness (*col9a1a, col9a2, col2a1a, col2a1b, col11a1a, col11a2*) and of genes for which the mouse orthologs have been implicated in hearing and/or ear development (*epyc, plod3, hapln1b, otos, clec3a, fgfr3, sox9a, acana, mycb, cdkn1bb, krtcap2*) (Supplemental Table 1). The combination of enriched expression of deafness and ear related genes that includes *sox9a* and many extracellular matrix proteins suggest that cluster 28 includes periotic mesenchyme cells. To begin to characterize cell-states within this putative periotic mesenchyme population, we further partitioned cluster 28 using the same unsupervised clustering pipeline into 8 different clusters, calculated their distance relationships, and then plotted the PAGA-initialized embedding (Fig. 6A and Supplemental Table 7) and coarse-grained PAGA-graphs (Fig. 6B). Again, the organization of cells in the PAGA-graph reflected their position in developmental time (Fig. 6C). The path begins with cell states in clusters m0-4 characterized by the expression of markers (*sox9b, foxc1b, serpinh1b*, subset of markers in Fig. 6D, markers for all clusters of Fig. 6A in Supplemental Table 7). Cluster m6 is characterized by expression of markers *clec3a*, epyc, and *otos-* all of which have been implicated in hearing in mouse models. A heatmap of marker gene expression and distance in gene expression space summarizes paths from immature precursors to cells of the periotic mesenchyme (Fig. 6E). We used HCR to visualize the expression of identified markers *otos, sox9b, foxc1b,* and *clec3a* in relationship to epithelial marker *epcam* (Fig. 6F). Other populations of putative periotic mesenchyme cells include clusters x1 and x2 that are characterized by the expression of marker *ndt5* and *f13a1b*.

**Fig. 6.**
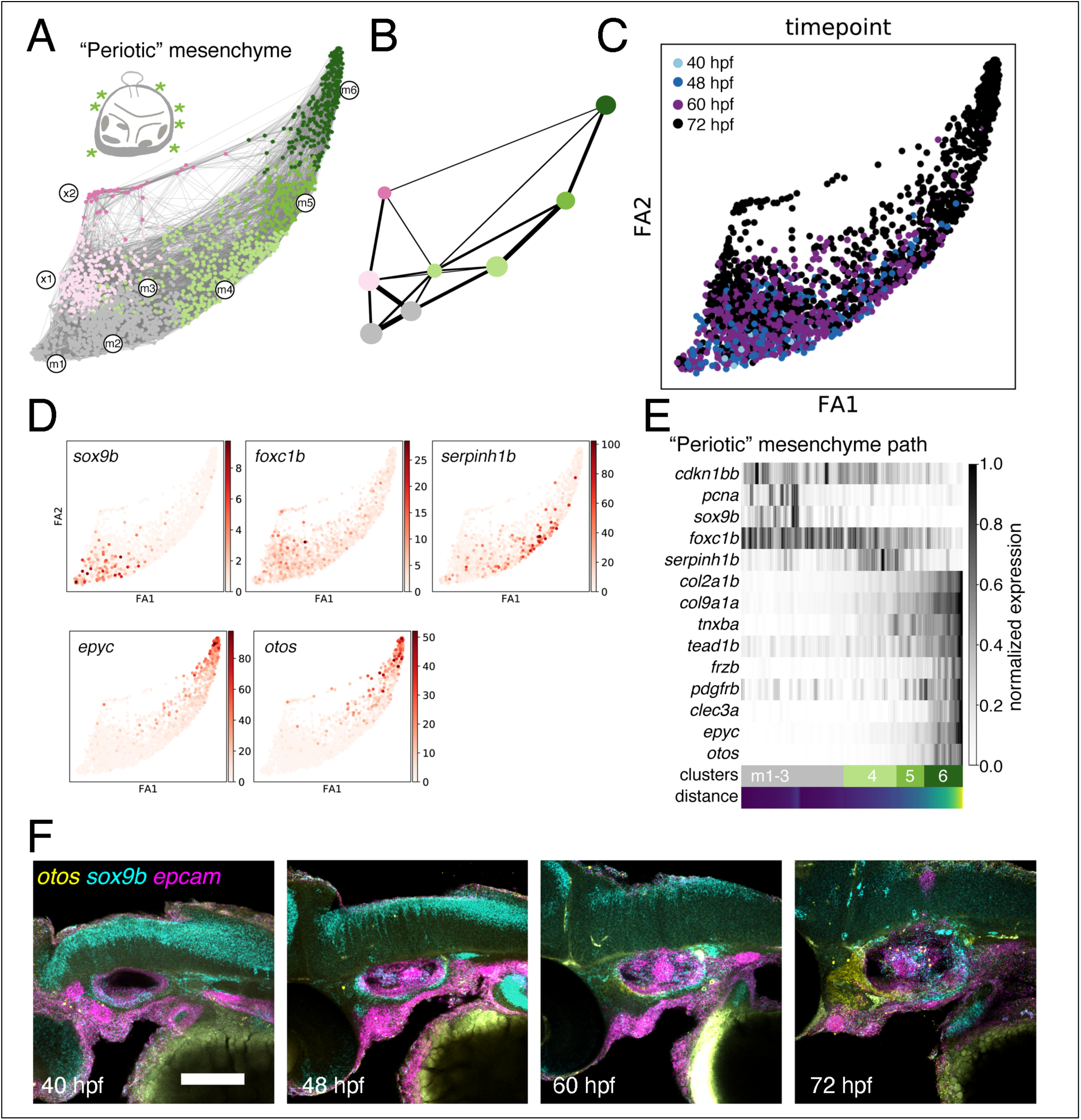
Fine analysis for single-cell transcriptomes of periotic mesenchyme cells. A. Cartoon schematic of a 3 day old zebrafish ear, with the periotic mesenchyme highlighted in green next to PAGA organized graph of periotic mesenchyme cells with more “mature” mesenchyme cells colored green. B. Coarse-grained PAGA graph of mesenchyme cells. C. Color-coding of developmental stage on PAGA graph from A. D. Heat-map of transcription counts on PAGA graphs with markers representative of different clusters. E. Collapsed heat-map histogram that compares gene expression in cells along the PAGA path with calculated distance along the path. F. Multiplexed fluorescent in situ analysis of periotic mesenchyme genes expressed at the queried developmental stages. (Scale bar 50 μm)

### *epcam* transcript levels are elevated in the endolymphatic sac of lmx1bb mutants

In addition to the analysis of the wild-type zebrafish ear, we also analyzed the single cell transcriptomes of ear enriched populations of cells from *lmx1bb* mutants (Fig. 7A), whose ES fails to relieve hydrostatic pressure(Swinburne et al. 2018). We identified differences between endolymphatic sac and duct cells that will provide an entry point for questions related to valve patterning (Supplemental Table 8). *Epcam,* encoding a regulator of cell adhesion molecules, is normally downregulated in the ES but this fails to happen in *lmx1bb* mutants (Fig. 7B-D). Epcam has been shown to inhibit lysosomal degradation of Claudin-7, which is expressed in the ES(Wu et al. 2013). Regulation of adhesion in the ES would provide opportunities for valve development, valve physiology, and valve mis-regulation in disease.

**Fig. 7.**
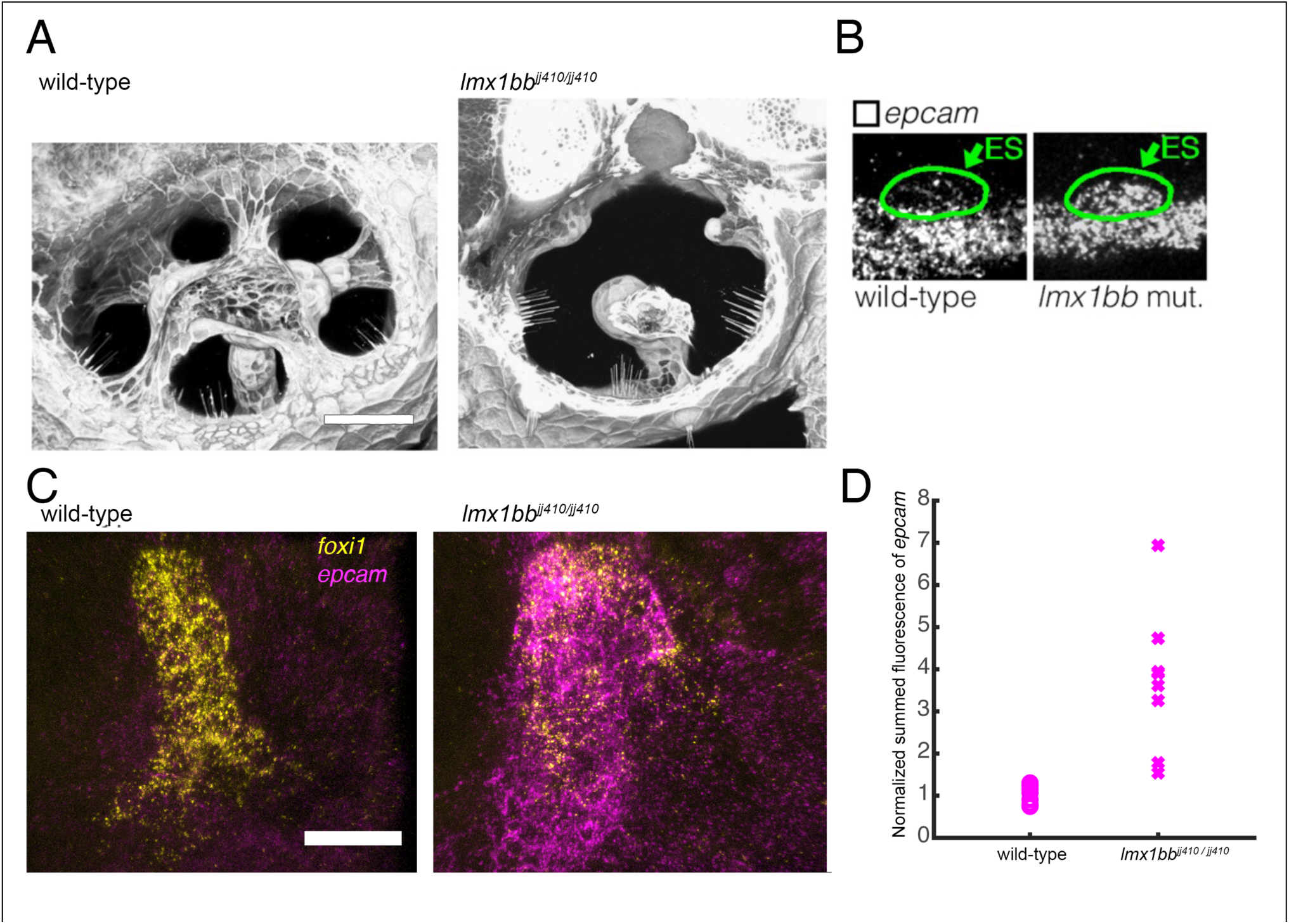
Comparison of wild-type vs *lmx1bb* mutant. A. 3-d rendering of confocal images of the wild-type and *lmx1bb* mutant ear at 3 dpf. B. A sagittal perspective of multiplexed fluorescent in situ analysis of *epcam* expression at 3 dpf. C. Maximum intensity projection of multiplexed fluorescent in situ analysis of *foxi1* and *epcam* 3 dpf. D. Quantification of in situs represented in C. (Scale bar 50 μm in A, 10 μm in C).

## Discussion

The embryonic ear forms by modification of cell states within lineages and positioning of nascent cell types in the correct place at the correct time. Molecular understanding of this system is critical for insights into human disorders of hearing or balance. Moreover, ear development is an ideal context in which to uncover general principles regarding how information flows between biochemical, mechanical, and geometric modes for vertebrate tissues (Collinet and Lecuit 2021). While many genes have been known to be important for ear development, to make further inroads into understanding the molecular networks whereby information flow requires a more complete parts list for the numerous different cell-states and behaviors involved. Here, we used scRNA-seq to assemble a more complete parts list that complements existing atlases of the zebrafish neuromasts and inner ear cells of other vertebrates (Honda et al. 2017; Janesick et al. 2021; Lush et al. 2019; Shrestha et al. 2018). Mining these resources opens numerous hypotheses and reveals unexpected molecular handles for unsolved mysteries of ear formation and function.

Studies of hair cell and support cell transcriptomes have enhanced our understanding of hair cell physiology and have attempted to determine why some organisms retain the ability to regenerate their hair cells while we cannot (Burns et al. 2015; Janesick et al. 2021; Lush et al. 2019; Shi et al. 2023). By comparing the hair cells of neuromasts to those of the otic vesicle, we have found evidence of parallel gene expression programs where different sources of hair cells often rely on different gene paralogs (*pvalb8* and *pvalb9*, *tmc2b* and *tmc2a, lhfpl5b* and *lhfpl5a,* Fig. 2G). Likewise, we uncovered clear differences between the two cell-types. For instance, NM hair cells differ from OV hair cells in their expression of genes implicated in stress responses such as *nqo1, gstp1,* and *nfe212a* (Fig. 2E).

Morphogenesis of SCCs requires differential expression of genes involved in the synthesis of a specialized ECM in the canal genesis zone. Yet the upstream developmental factors instructing this pattern have remained unknown. Our dataset suggests *ccn1l1* (cell communication factor 1-like 1) as a likely candidate for this process. The protein product of *ccn1l1* (also known as *cyr61l1)* is predicted to be secreted in the extracellular region and to have heparan sulfate binding activity resulting in signal transduction (Babic et al. 1998; Chaqour and Goppelt-Struebe 2006; Mugahid et al. 2020; Si et al. 2006). These properties make it an excellent candidate morphogenic factor, whose functional characterization will reveal roles not only in canal morphogenesis but also in other embryonic tissues where it is expressed.

Advances in understanding the development and functional role of the endolymphatic duct and sac have become more accessible in the zebrafish system, where the in vivo tissues are optically accessible when the ear is functional. In the past, we reported that the endolymphatic sac can behave as a tissue-scale relief valve (Swinburne et al. 2018). Here, we identified numerous molecular handles for understanding how this tissue behavior develops and begins to work. We found that the contractile protein Smoothelin is specifically expressed in the distal tip of the ES. We also uncovered a potentially important effector of Lmx1b activity in the endolymphatic sac. Specifically, loss of Lmx1b leads to elevated expression of *epcam* in the ES, a known regulator of cell-cell adhesion. In addition, we found that loss of Lmx1b can prevent the expression of *smoothelin* in the ES. These findings indicate that the Lmx1b control system in the ES can both positively and negatively regulated gene expression. Future studies will explore the roles of these and other distinctive ES genes that we have uncovered.

Historically, the relationship between the ear anlage and its associated mesenchyme has intrigued embryologists as a system of developmental cross-induction (Kaan 1930, 1938; McPhee and Van de Water 1986). Molecular studies have uncovered roles for retinoic acid, TGF-B, FGF, SHH, Wnt, and BMP signaling in this inductive back-and-forth (Billings et al. 2021; Frenz and Liu 2000; Ladher et al. 2005; W. Liu et al. 2002; Wei Liu et al. 2003, 2008). In addition to identifying a population of periotic mesenchymal cells expressing numerous deafness genes, analysis of these cells’ transcriptomes identified many genes encoding ligands or receptors known to be involved in cell-cell signaling during various development processes—ligands included *bmp1a, bmp2b, igf2b, tgfbi, wnt11,* and *notch2* and receptors included *fgfr2*, *fgfr4, fgfr3, pdgfrb, fzd2,* and *fzd7b* (Supplemental Table 7). Knowing the identities of these signaling factors will help decipher how the periotic mesenchyme communicates with the developing ear for cross-induction.

The gene-lists of developmental scRNA-seq atlases help distinguish cell-states or cell-types, especially rare ones, that may hold critical importance for a process or organ physiology. Nevertheless, our approach lacks lineage analysis of our inferred cell-state relationships and future studies will require cell recording technology via molecular barcodes and lineage analysis via live-imaging (Kester and van Oudenaarden 2018; McKenna et al. 2016; Wagner and Klein 2020; Wagner et al. 2018).

In recent years, there has been a significant increase in the number of single-cell datasets available for studying embryonic development, providing a crucial foundation for important mechanistic studies aimed at understanding fundamental biological questions(Sur et al. 2023). Our single-cell transcriptomic atlas of ear morphogenesis is a resource for uncovering the molecular control networks that contribute to the development of the ear’s hair cells, semicircular canals, endolymphatic duct and sac, and the periotic mesenchyme. Molecular handles uncovered here lay the groundwork for uncovering mechanisms by which: the ear’s sensory cells differ from the neuromast’s sensory cells, the semicircular canals are positioned, the endolymphatic sac promotes ear homeostasis, and how the periotic mesenchyme contributes to ear morphogenesis and how the ear promotes development of the temporal bone.

## Materials and Methods

### Zebrafish strains and maintenance

Zebrafish were maintained at 28.5°C using established protocols (Westerfield 1993). The Harvard Medical Area Standing Committee on Animals approved zebrafish work under protocol number 04487 and the University of California, Berkeley zebrafish work was approved by UCB animal protocol #AU 2020-10-13737. Adult zebrafish, 3 months to 2 years of age, were mated to produce embryos and larvae. Adults were housed in a main fish facility containing a 19-rack Aquaneering system (San Diego, CA) and a 5-rack quarantine facility for strains entering the system from outside labs or stock centers. The systems’ lights turn on at 9 am and go off at 11 pm. Fish matings were set up the night before with males separated from females with a divider in false-bottom crossing cages in a pair wise, or 2 × 2 fashion to maximize embryo yield. The divider was pulled the following morning, shortly after the lights turned on. Egg production was monitored to establish the time of fertilization. Manipulations and observations of baby zebrafish were performed between 40 and 72 hours post fertilization. These studies used AB wild-type and *lmx1bb^jj410^* strains (Schibler and Malicki 2007). *Lmx1bb ^jj410^* mutants were maintained as heterozygous adults with additional mutations that minimize pigmentation: *mpv17^a9^*, *mitfa^w2^*, and *csfr1a^j4e1^* (Parichy et al. 2000; White et al. 2008). This pigment free triple mutant was used as the background for all fluorescent in situ analyses of wild-type and *lmx1bb^jj410^* embryos.

### Dissections and tissue dissociation

To minimize sticking of cells to plastic, 1.5 ml collection tubes were coated with 10% BSA for 10 minutes, rinsed in PBS, and left on ice. Dissections were performed in modified DMEM plus tricaine: 1x DMEM (11039-021 (LifeTech)), 10 mM HEPES, 1x non-essential amino acids (11140076 (LifeTech)), 1mM sodium pyruvate (11360070 (LifeTech), 100 um B-mercaptoethanol (M3148 Sigma), and 0.33 mg/ml Tricaine (E10521 (Sigma)). The morning of an inDrops experiment, a team of 3-4 researchers dissected otic vesicle enriched tissue as in Fig. 1D using two 22-gauge trabecular needles with syringes for handles. For each inDrops run, 40-50 embryos were dissected, and the tissue was collected in a coated collection tube. The DMEM was removed and 50 µl of Trypsin-EDTA was added. Avoiding air bubbles, the tissue was pipetted up and down several times with a 200 µl pipette tip. The tissue was left on ice to dissociate for 3 minutes for 36 hpf tissue, 5 minutes for 48 hpf tissue, or 7 minutes for 60 hpf tissue. We then added 100 µl of FACSMax, pipetted the sample up and down (avoiding air bubbles) and incubated on ice for 2 minutes. While incubating, we rinsed out a BSA coated tube with PBS. We then added 850 µl of FACSmax to the dissociated cells and pipetted through a 40 µm cell filter into a rinsed BSA coated tube. The samples were spun for 3 minutes at 300g in a cooled swinging bucket centrifuge. Supernatant was removed and cells were resuspended in 1 ml of 0.5% BSA in chilled PBS. We then spun down the cells again for 3 minutes at 300g in a cooled swinging bucket centrifuge. Supernatant was removed and cells were resuspended in 200 µl of 1x prechilled PBS. Cells were counted using a hemocytometer, diluted to 800,000 cells/ml and then used to collect ∼15,000 cells by inDrops. Dissection, dissociation, and cell collection was performed twice for each condition.

### inDrops collection, library preparation, and analysis

Single-cells were captured and transcriptome libraries were prepared by the single-cell core at Harvard Medical School with established protocols (Klein et al. 2015; Wagner et al. 2018). Libraries were pooled in a manner to evenly distribute samples and prevent overlapping of barcodes. Assembled libraries were then sequenced with paired end reads using 19 flow cells of 75 cycles on an Illumina NextSeq 500. As described previously, cDNA reads were mapped using the GRCz11 zebrafish genome assembly (Wagner et al. 2018).

Data analysis was performed using SCANPY and implementations of Jupyter notebooks are available (https://github.com/orgs/SwinburneLabUCB/repositories) (Kluyver, et al. 2016; Wolf et al. 2018). Depending on the quality of each sample determined by barcode counts histograms, cells were removed if they had fewer than 250-1200 UMIs. Cells with greater than 15% UMIs coming from mitochondrial genes were also removed. The counts for each cell’s transcriptome were then normalized to the mean of that cell for all genes and genes were removed that were present in fewer than 3 cells. All data was pooled into a single AnnData library and unique indices were added to each cell, “cell_ix”, to simplify reanalysis of cell subsets. To reduce the dimensionality and cluster similar cell states, counts per cell were normalized and highly variable genes were identified based on expression in more than 3 cells, a minimum mean of 0.01, a maximum mean of 3, and minimum dispersion of 0.5. Cell cycle and housekeeping-correlated genes were identified and then excluded by seeding a list of previously identified cell cycle and housekeeping genes and expanding the list to include genes in our cell population with correlation coefficients greater than 0.4 (see ‘Full_Filter_cm_v1.csv’) (Tirosh et al. 2016; Wagner et al. 2018). Principle component analysis was performed to identify a lower dimensional space on which to project the normalized gene expression of the highly variable genes (Pedregosa et al., n.d.). A neighborhood graph was then computed and used to embed the graph using UMAP (McInnes et al. 2020). The graph was clustered into subgroups using the Leiden algorithm (Traag et al. 2019). For each Leiden cluster, marker genes were identified using a Wilcoxon rank-sum test implemented with a false discovery rate correction using the Benjamin-Hochberg procedure. For clusters of interest, the list of cell indices was recorded, and the subset of cells was reanalyzed. PAGA analysis, partition-based graph abstraction, was implemented to generate more interpretable maps of cell positions in gene expression space (Wolf et al. 2019).

### HCR fluorescent in situ hybridization

Fluorescent in situ hybridization by hybridization chain reaction (HCR, Molecular Instruments) experiments were performed on whole mount embryos fixed at 40, 48, 60, or 72 hpf as previously described (Choi et al. 2018b). Embryos were fixed in 4% paraformaldehyde (PFA) and then stored in methanol at −20°C. Fixed embryos were rehydrated in a series of methanol and PBST (0.1% Tween) washes and permeabilized with 10 ug/ml proteinase K (10 minutes for 40 hpf, 20 minutes for 48 hpf, and 25 minutes for 72 hpf). Embryos were then refixed in 4% PFA and then prehybridized in probe hybridization buffer at 37°C. Pools of gene specific probes were then added at a final concentration of 4nM and incubated overnight in a 37°C hybridization oven with mixing on a rotisserie. On the following day, embryos were washed with 4 times with probe wash buffer at 37°C and 2 time with 5x SSCT (5x SSC with 0.1% Tween 20) at room temperature. Embryos were then preincubated in amplification buffer as the appropriate amplification hairpins were prepared. Hairpins were added and amplification was performed overnight at room temperature with mixing. On the following day, samples were washed with SSCT and then imaged or stored at 4°C. To counteract shrinkage artifacts when preparing the semicircular canal samples (Fig. 4), we adapted an acid fixation method to protonate molecules within samples to promote the retention of water (Fuentes and Fernández 2014). Prior to HCR, embryos were transferred to 3 ml of 4% paraformaldehyde and 4 drops of glacial acetic acid (∼200 ul) were added and mixed with the sample. The acid fixation samples were fixed for 2 hours at room temperature with mixing and then entered a standard HCR protocol after the acid fixation. For acid fixed samples, proteinase K digestion times were reduced by 50%.

Images of HCR samples were collected on an upright LSM 710 (Carl ZEISS, Göttingen, Germany) or an upright LSM 980 (Zeiss) with a c-aprochromat 40x/NA 1.2 objective (Zeiss) or a 20x/NA 1 objective (Zeiss). Samples were mounted using custom agarose molds as described previously (Swinburne et al. 2018). Lasers of wavelength 488, 561, and 639 nm were used to excite samples developed with HCR amplifiers labelled with AlexaFluor 488, 546, and 647, respectively. We quantified and locally normalized the summed fluorescence of signals in the endolymphatic duct and sac using the *foxi1* pattern to manually segment the tissue with Fiji software (Schindelin et al. 2012).

## Acknowledgements

We thank Andrew Murphy for zebrafish care at Harvard Medical School. We thank Tyler Mentley, Kait Kliman, Leah Ferger, Ian Strieter, Sandra Wiese, Frances Campbell, Lindsey Arenson, and the rest of the Aquatics Facility Team for zebrafish care at UC Berkeley. We thank Mandovi Chatterjee and the rest of the Single Cell Core at Harvard Medical School for single cell isolation and library preparation. We thank Sandy Nandagopol for sharing a python script for generating HCR probe-sets. This work was supported by Hearing Health Foundation Emerging Research and Meniere’s Disease Grants (IS), R01 DC021710 from the National Institute of Deafness and Other Communication Disorders (IS), and K99 HD098918 (AM). This work was supported by R01 DC010791 and R01 DC015478 from the National Institute of Deafness and Other Communication Disorders (SGM).

