## Supplementary figures and images for "A single-cell transcriptomic atlas of inner ear morphogenesis in zebrafish"

### Supplemental Figure 1

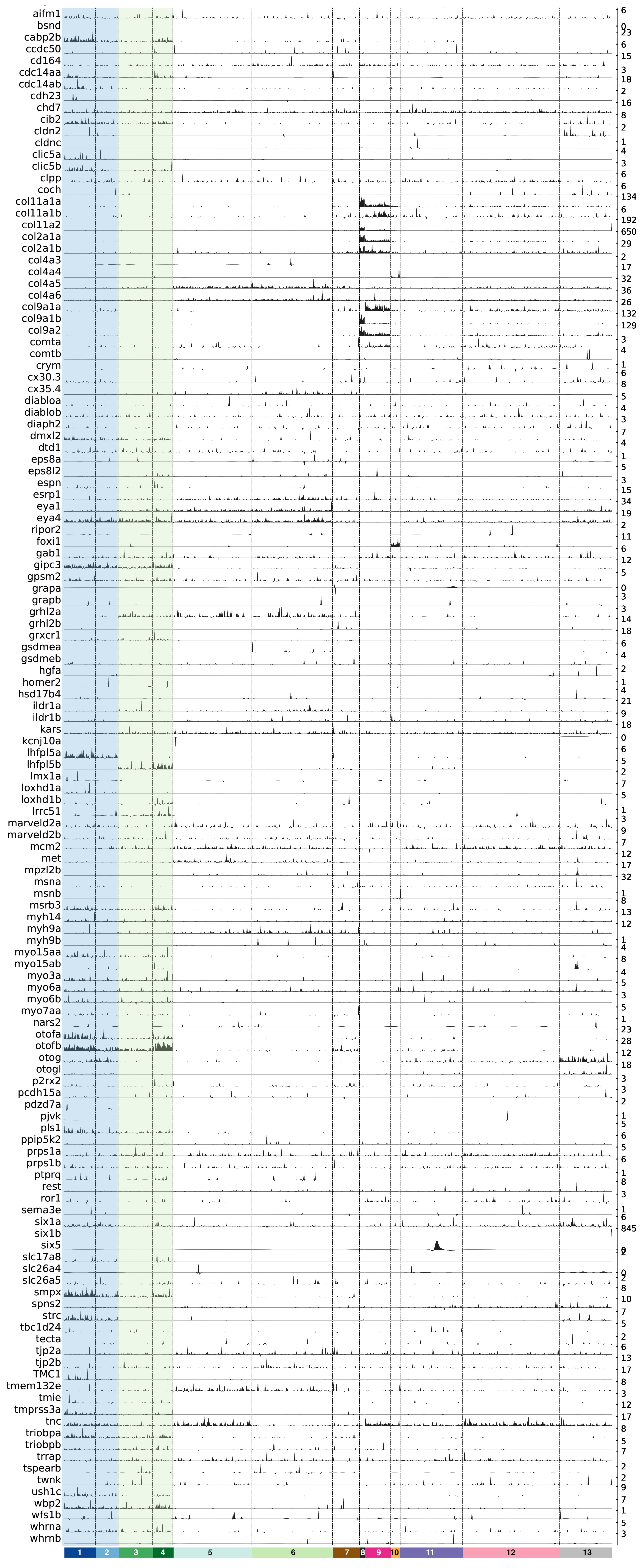
